# Making Biomedical Research Software FAIR: Actionable Step-by-step Guidelines with a User-support Tool

**DOI:** 10.1101/2022.04.18.488694

**Authors:** Bhavesh Patel, Sanjay Soundarajan, Hervé Ménager, Zicheng Hu

## Abstract

Findable, Accessible, Interoperable, and Reusable (FAIR) guiding principles tailored for research software have been proposed by the FAIR for Research Software (FAIR4RS) Working Group. They provide a foundation for optimizing the reuse of research software. The FAIR4RS principles are, however, aspirational and do not provide practical instructions to the researchers. To fill this gap, we propose in this work the first actionable step-by-step guidelines for biomedical researchers to make their research software compliant with the FAIR4RS principles. We designate them as the FAIR Biomedical Research Software (FAIR-BioRS) guidelines. Our process for developing these guidelines, presented here, is based on an in-depth study of the FAIR4RS principles and a thorough review of current practices in the field. To support researchers, we have also developed a workflow that streamlines the process of implementing these guidelines. This workflow is incorporated in FAIRshare, a free and open-source software application aimed at simplifying the curation and sharing of FAIR biomedical data and software through user-friendly interfaces and automation. Details about this tool are also presented.

## Introduction

Research software (including scripts, computational models, notebooks, code libraries, etc.) has become an increasingly important part of scientific research. A survey conducted by the Software Sustainability Institute in the UK has found that 92% of academics use research software, 69% say that their research would not be practical without it, and 56% develop their own software^1^. Other surveys have similarly shown the importance of research software in scientific research^2–4^. Research software plays a fundamental role not only in collecting, analyzing, and processing data but has also become the centerpiece of many scientific research projects aimed at developing computational models to understand and predict various physical phenomena. In line with this general trend, research software has also become an essential part of biomedical research over the last decade, especially with the advent of machine learning and artificial intelligence in the field. The evolution of the number of new biomedical-related software repositories created on GitHub every year, shown in **Figure 1**, gives an overview of this trend.

**Figure 1.**
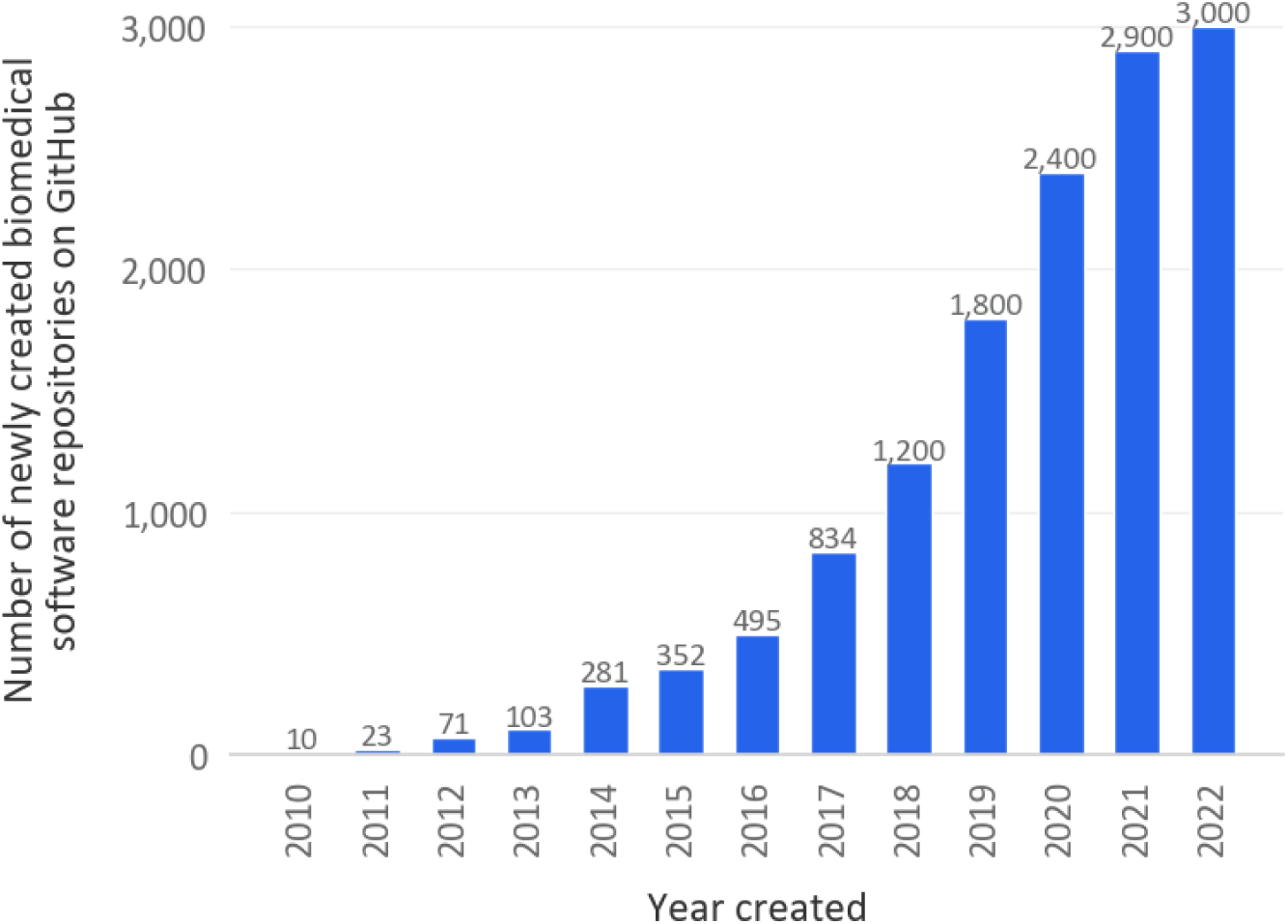
Evolution of the number of new biomedical-related software repositories created on GitHub in a given year (rounded to the hundredth when >1,000). The search consisted of looking for new repositories created in a given year with “biomedical” included in their name, tags, README, or description. To exclude repositories with data only (e.g., CSV files or markdown text), the search was limited to repositories with a major coding language being one of the popular software programming languages. This list of popular software programming languages was established based on GitHub’s list of popular programming languages to which we added a couple of languages that we deemed relevant for biomedical research software. For more details, see the code associated with this manuscript (c.f. Code Availability section).

Research software has consequently become an essential asset of scientific research and it has therefore become critical to preserve, share, and make it reusable. The Findable, Accessible, Interoperable, and Reusable (FAIR) guiding principles provide a foundation for achieving that^5^. Published in 2016, these principles are aimed at optimizing data reuse by humans and machines. While postulated for all digital research objects, several research groups have shown that the FAIR principles as written do not directly apply to software because they do not capture the specific traits of research software^6, 7^. In 2019, Lamprecht et al. were the first to propose reformulated FAIR principles that are tailored to research software. They were published in 2020^6^. Noticing yet a need for more software-specific principles, the FAIR for Research Software (FAIR4RS) Working Group, jointly convened as a Research Data Alliance (RDA) Working Group, FORCE11 Working Group, and Research Software Alliance (ReSA) Task Force, initiated a large-scale effort shortly after in mid-2020 to tackle this need. A subgroup of this working group took a fresh look at the FAIR principles and their applicability to research software, and suggested a set of reformulated FAIR principles tailored for research software that were published as a preprint in January 2021^8^ and then as a peer-reviewed article in March 2021^9^. Based on that effort and the efforts of other subgroups, the FAIR4RS working group published in June 2021 a draft for formal community review of their new principles called the FAIR4RS principles (v.03)^10^. After revising the FAIR4RS principles based on community feedback, they published the final version (v.1.0) in May 2022^11^. This version was introduced in a peer-reviewed article in September 2022^12^.

Just like the original FAIR principles, the FAIR4RS principles are aspirational and aimed at providing a general framework. One study provides some actionable guidelines for making research software FAIR but it does not include a clear process and is based on the original FAIR principles rather than the FAIR4RS principles^13^. Another study also provides some actionable guidelines for making research software FAIR but it is not aimed at complying with each principle and is also based on the original FAIR principles^14^. Actionable guidelines that researchers can follow for complying with each of the FAIR4RS principles are not yet available, which is hindering the widespread adoption of FAIR practices. Particularly in biomedical research, the COVID-19 pandemic has emphasized the need for such guidelines^15^. As noted by the RDA COVID-19 Working Group: “Whilst preprints and papers are increasingly openly shared to accelerate COVID-19 responses, the software and/or source code for these papers is often not cited and hard to find, making reproducibility of this research challenging, if not impossible”^15^.

To fill this gap, we proposed in this work the first minimal and actionable step-by-step guidelines for researchers to make their biomedical research software compliant with the FAIR4RS principles. Our focus was on biomedical research software given that this effort was initiated as part of a larger project aimed at supporting the curation and sharing of COVID-19 related research data and software. We designate these guidelines as the FAIR Biomedical Research Software (FAIR-BioRS) guidelines. Many challenges have been reported in establishing such actionable guidelines, such as the lack of community agreement on metadata and identifiers^8^. Rather than waiting for these challenges to be overcome, our approach consisted of deriving the best actionable guidelines possible around these challenges such that researchers can already make their research software FAIR. The FAIR-BioRS guidelines and our approach to deriving them are presented in this manuscript.

Given that funding to support the reusability of software is lacking, the main responsibility of making research software FAIR falls on the researchers developing the software, i.e., they may receive only limited support to assist them in this effort. Our proposed guidelines can thus add an additional burden on these researchers who are already facing significant challenges when developing their software^16–18^. To address this concern and ensure the FAIR-BioRS guidelines are easily accessible and widely embraced by researchers, we have developed a workflow to streamline the process of implementing the FAIR-BioRS guidelines. This workflow is incorporated in FAIRshare, a free and open-source desktop software application aimed at simplifying the processes for making biomedical data and software FAIR. FAIRshare takes the user’s software-related dataset as input and walks them step-by-step into implementing the FAIR-BioRS guidelines. FAIRshare provides an intuitive graphical user interface at each step where the user can easily provide required inputs for items that cannot be automatically assessed (e.g., selecting a desired license for the software), and includes automation in the backend to take over complex and/or time-consuming tasks that can be automated (e.g., creating a LICENSE file with standard license terms once the license is selected). Details about this tool are also provided in this manuscript.

## Results

### Overview

The work in this manuscript is based on the FAIR4RS principles v1.0^11^ given that they are the most thorough to date and backed by a large community support. Our work was initiated in December 2021 and was initially based on the formal draft version of the FAIR4RS principles (v.0.3)^10^. We then reiterated on our work between November 2022 and June 2023 to align with the final version (v1.0) of the FAIR4RS principles as well as incorporate community feedback we received along the way. Our overall process for establishing the FAIR-BioRS guidelines is illustrated in **Figure 2**. More details are provided in the Methods section.

**Figure 2.**
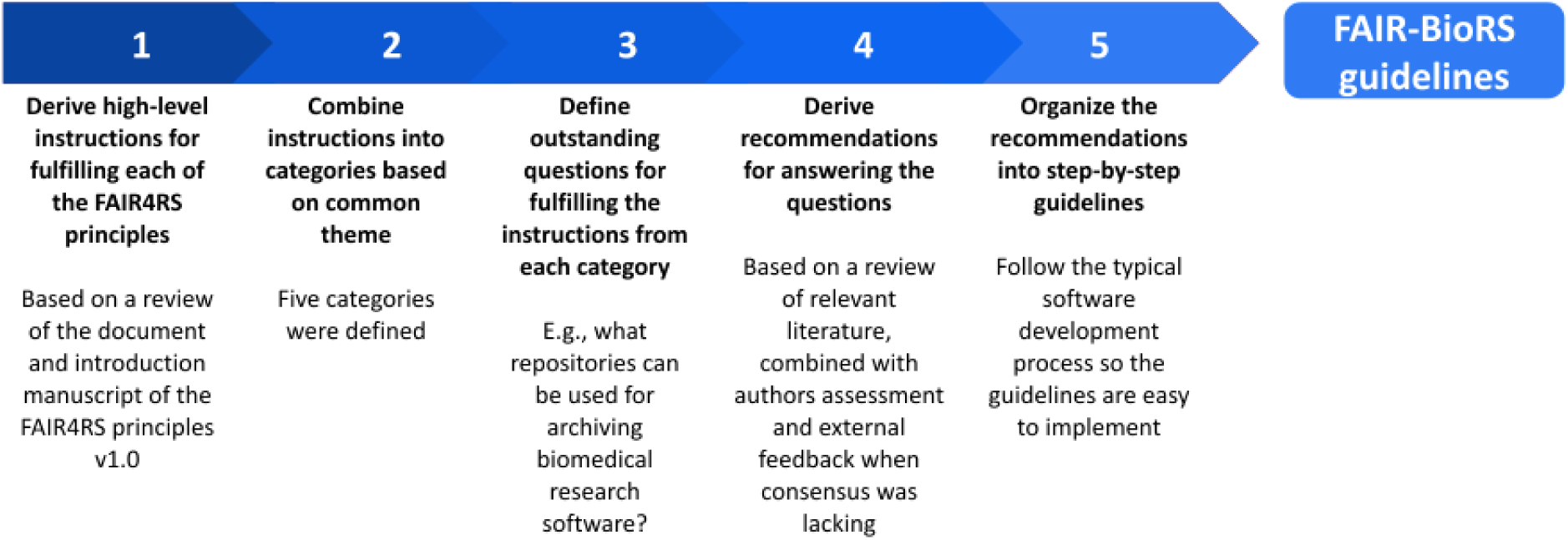
Overall process followed to establish the current version of the FAIR-BioRS guidelines.

### High-level categories and instructions to fulfill the FAIR4RS principles

For each of the FAIR4RS principles, we derived concise high-level instructions to fulfill that principle based on related information available in the FAIR4RS v1.0 publication^11^ and the associated introduction manuscript^12^. These instructions are available in the “fair4rsv1.0Instructions” tab of the “data.xlsx” file associated with this manuscript (c.f. Data Availability section). We identified that the instructions for the different principles had overlapping themes and therefore organized the instructions into five action-based categories to help in our process of deriving actionable guidelines:

- Category 1: Develop software following standards and best practices
- Category 2: Include metadata
- Category 3: Provide a license
- Category 4: Share software in a repository
- Category 5: Register in a registry

The instructions associated with each category and the FAIR4RS principles they cover are provided in **Table 1**. Then, for each category, we identified outstanding questions we needed to answer for deriving actionable guidelines that allow us to fulfill the high-level instructions from that category. These questions are also provided in **Table 1**. To answer these questions, we combined findings from a review of relevant resources, our own understanding of research software development, and external suggestions we received from various communities (c.f. Discussion section) to derive relevant recommendations. This is presented next.

**Table 1.**
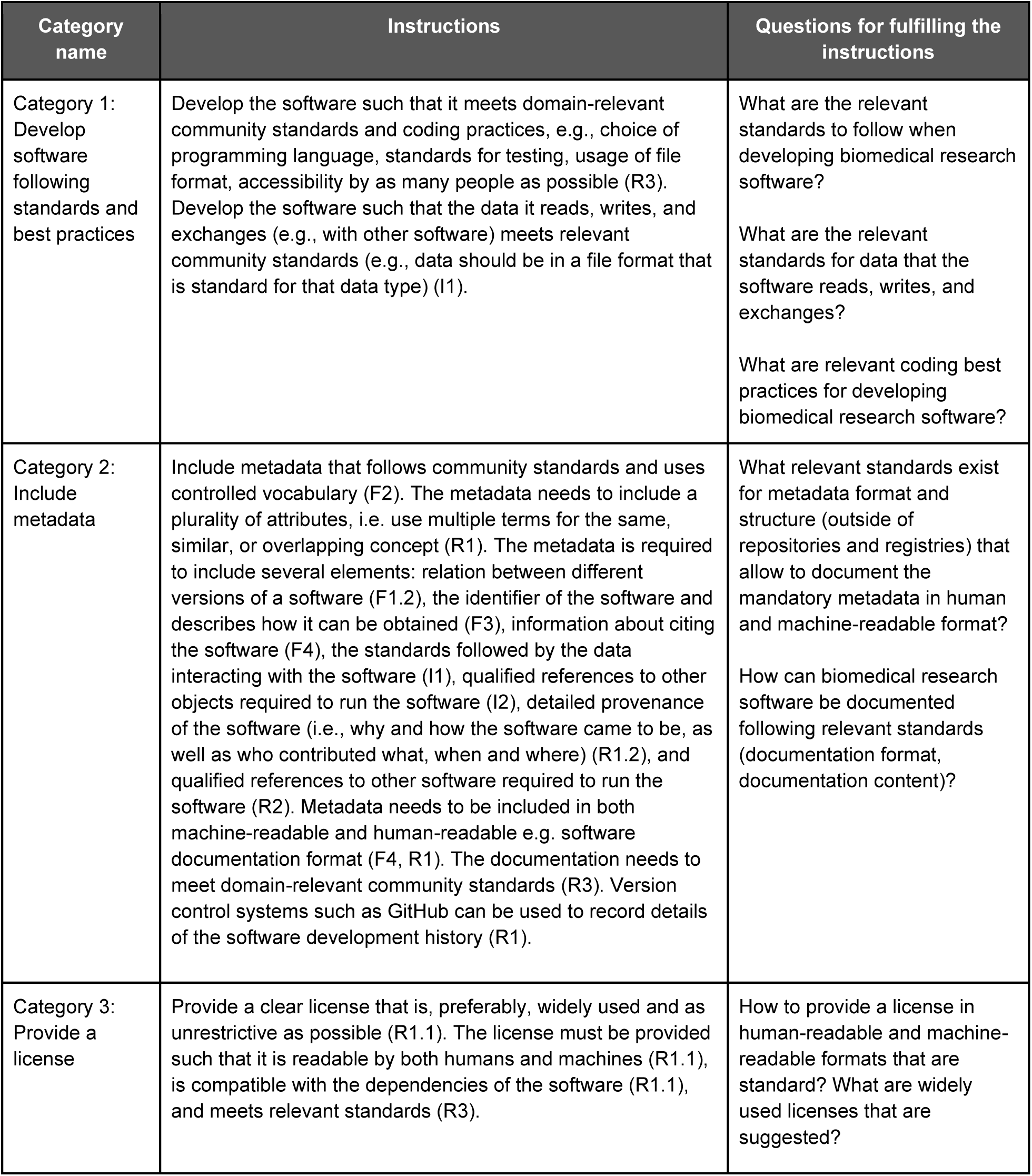
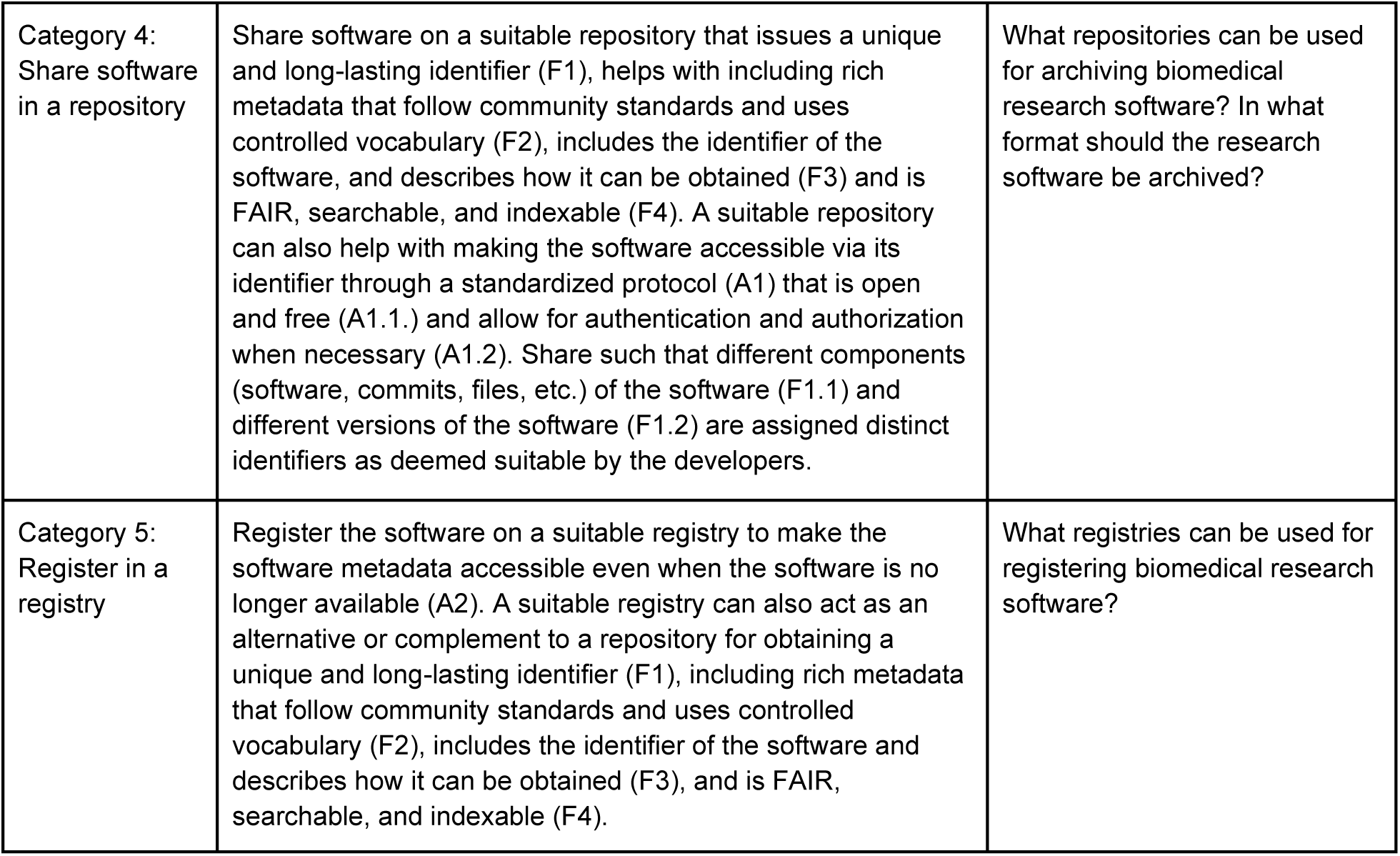
The five categories were established based on the instructions we derived for each of the FAIR4RS principles v1.0. The combined instructions for each category, the FAIR4RS principles they cover, and the outstanding questions for fulfilling these instructions are also provided.

| Category name | Instructions | Questions for fulfilling the instructions |
| --- | --- | --- |
| Category 1:<br>Develop software following standards and best practices | Develop the software such that it meets domain-relevant community standards and coding practices, e.g., choice of programming language, standards for testing, usage of file format, accessibility by as many people as possible (R3). Develop the software such that the data it reads, writes, and exchanges (e.g., with other software) meets relevant community standards (e.g., data should be in a file format that is standard for that data type) (I1). | What are the relevant standards to follow when developing biomedical research software?<br>What are the relevant standards for data that the software reads, writes, and exchanges?<br>What are relevant coding best practices for developing biomedical research software? |
| Category 2:<br>Include metadata | Include metadata that follows community standards and uses controlled vocabulary (F2). The metadata needs to include a plurality of attributes, i.e. use multiple terms for the same, similar, or overlapping concept (R1). The metadata is required to include several elements: relation between different versions of a software (F1.2), the identifier of the software and describes how it can be obtained (F3), information about citing the software (F4), the standards followed by the data interacting with the software (I1), qualified references to other objects required to run the software (I2), detailed provenance of the software (i.e., why and how the software came to be, as well as who contributed what, when and where) (R1.2), and qualified references to other software required to run the software (R2). Metadata needs to be included in both machine-readable and human-readable e.g. software documentation format (F4, R1). The documentation needs to meet domain-relevant community standards (R3). Version control systems such as GitHub can be used to record details of the software development history (R1). | What relevant standards exist for metadata format and structure (outside of repositories and registries) that allow to document the mandatory metadata in human and machine-readable format?<br>How can biomedical research software be documented following relevant standards (documentation format, documentation content)? |
| Category 3:<br>Provide a license | Provide a clear license that is, preferably, widely used and as unrestrictive as possible (R1.1). The license must be provided such that it is readable by both humans and machines (R1.1), is compatible with the dependencies of the software (R1.1), and meets relevant standards (R3). | How to provide a license in human-readable and machine-readable formats that are standard? What are widely used licenses that are suggested? |
| Category 4:<br>Share software in a repository | Share software on a suitable repository that issues a unique and long-lasting identifier (F1), helps with including rich metadata that follow community standards and uses controlled vocabulary (F2), includes the identifier of the software, and describes how it can be obtained (F3) and is FAIR, searchable, and indexable (F4). A suitable repository can also help with making the software accessible via its identifier through a standardized protocol (A1) that is open and free (A1.1.) and allow for authentication and authorization when necessary (A1.2). Share such that different components (software, commits, files, etc.) of the software (F1.1) and different versions of the software (F1.2) are assigned distinct identifiers as deemed suitable by the developers. | What repositories can be used for archiving biomedical research software? In what format should the research software be archived? |
| Category 5:<br>Register in a registry | Register the software on a suitable registry to make the software metadata accessible even when the software is no longer available (A2). A suitable registry can also act as an alternative or complement to a repository for obtaining a unique and long-lasting identifier (F1), including rich metadata that follow community standards and uses controlled vocabulary (F2), includes the identifier of the software and describes how it can be obtained (F3), and is FAIR, searchable, and indexable (F4). | What registries can be used for registering biomedical research software? |

### Review of current practices and resulting recommendations

#### Reviewed studies

Our review approach is presented in the Methods section. A list of the reviewed resources is available in the “resourcesList” tab of the “data.xlsx” file included in the dataset associated with this manuscript (c.f. Data Availability section). In these resources, we looked for actionable recommendations that answer any of the questions listed in **Table 1**. A total of 39 resources were deemed relevant to these questions and included in the analysis presented next. Details about the information collected from each resource are provided in the “resourcesReview” tab of the “data.xlsx” file included in the dataset associated with this manuscript (c.f. Data Availability section). We summarize below key findings for each of the categories and provide our resulting recommendations.

#### Category 1: Develop software following standards and best practices

None of the reviewed resources provided actionable items for following standard development practices, except one that mentions the PEP 8 Style Guide for Python Code (http://peps.python.org/pep-0008). This is understandable since such standards vary depending on many factors such as the domain of research and coding language.

Similarly, there are no clear standards provided for the format of the data that software read, write, and exchange. Few of the reviewed resources^10, 11, 19^ refers to the FAIRsharing Registry (https://fairsharing.org) for a curated list of community standards^20^.

Suggestions for best development practices are provided in 21 of the reviewed resources. Some of the reviewed resources are fully dedicated to best practices, including general best practices for scientific software development^21^ and specific best practices for biomedical software^22, 23^. Overall, developing with a version control system is commonly suggested^8, 12–15, 19, 21–28^. GitHub, BitBucket, and GitLab are commonly mentioned ready-to-use version control system platforms. Using container technologies such as Docker and Singularity^13, 18, 19, 22, 24, 29^, having code level documentation (in code comments, description in the headers)^19, 21, 22, 25, 30^, and recording dependencies^11, 12, 19, 21^ are other best practices mentioned in the reviewed resources.

There is clearly a lack of community-agreed standards and best practices. Therefore, we used findings from this review, combined with our own assessment and external suggestions we received when consensus was lacking, to establish our recommendations for fulfilling the instructions from Category 1. We recommend working from a version control system platform (e.g., GitHub, Bitbucket, GitLab) as this seems essential for complying with several of the FAIR4RS principles. We also recommend having code-level documentation (e.g., in code comments, description in the headers) when deemed necessary for code reusability. Additionally, we recommend recording dependencies as per standard practices for the coding language (e.g., in a requirement.txt file for Python code, in a package.json file for Node projects, or in a DESCRIPTION file for R packages). Some of the dependencies can also be recorded in the software documentation such as in the README file (see documentation discussion in Category 2 below). Following language-specific standards and best practices, which depend on the development stack used, is also recommended. For instance, we suggest following the PEP 8 Style Guide for Python Code or Google’s R Style Guide for R code (https://google.github.io/styleguide/Rguide). Finally, we recommend ensuring that inputs/outputs of the software follow any applicable community standards (e.g., General Feature Format (GFF) for genomic annotation files) as this is essential for interoperability. We refer to FAIRsharing Registry for finding relevant standards for the biomedical domain of interest in the software. We leave out recommendations on using container technologies as this is a complex topic that we deem out of the scope of the minimal guidelines we are aiming for here.

#### Category 2: Include metadata

From the reviewed resources, 24 have made a recommendation of metadata files and ontologies to use for research software. A majority of them^6, 8, 10, 12–14, 17–19, 24, 25, 31–36^ suggest following the guidelines of the CodeMeta Project for including metadata in research software. The CodeMeta Project is an academic-led community initiative that was the result of the FORCE11 Software Citation working group. It aims to formalize the metadata fields included in typical software metadata records and introduces important fields that did not have clear equivalents. The CodeMeta vocabulary resulting from this effort is built over the schema.org classes SoftwareApplication and SoftwareSourceCode, which links the data for semantic web discovery. Some additional terms necessary to describe software that are not part of schema.org have been included in the CodeMeta schema. There is a continued effort to propose these terms to schema.org so that the CodeMeta schema completely aligns with schema.org rather than becoming a separate standard. Metadata information conformant to the CodeMeta vocabulary is to be included in JSON-LD format in a file named codemeta.json and stored in the root directory of the software. The Software Heritage group has developed a CodeMeta generator (https://codemeta.github.io/codemeta-generator) that can be used to create a codemeta.json file or edit an existing one. Tools are also available to include a codemeta.json file in R packages^37^.

Including a citation file following the Citation File Format (CFF) is another popular suggestion amongst the reviewed resources^13, 14, 18, 19, 25–28, 30, 34–36^. The CFF was developed by a group of academics assembled under the group “Development and implementation of a standard format for CITATION files”^38^. The goal of the CFF is to provide an all-purpose citation format and specifically provide optimized means for the citation of software via the provision of software-specific reference keys and types. CFF files must be named CITATION.cff, implemented in YAML 1.2, which is a machine-readable format that optimizes human readability, and must be stored in the root directory of the software. The CFF file initializer (https://citation-file-format.github.io/cff-initializer-javascript) is mentioned by several reviewed resources as a tool for easily generating a CITATION.cff file. Note that the CITATION.cff file integrates with the GitHub citation feature such that if a CITATION.cff file is included in the root folder of a GitHub repository, a “Cite this repository” option is automatically displayed on the repository landing page making it easier to cite a software repository on GitHub.

The codemeta.json and CITATION.cff are general metadata files that can be used for any research software. One of the reviewed resources^6^ suggests including language-specific metadata for instance based on the DESCRIPTION file for R packages or the PEP 566 for Python packages. Other reviewed resources^6, 24, 25^ also suggested preparing metadata using bio-specific ontologies such as EDAM^19, 39, 40^, biotoolsSchema^41^, and Bioschemas^42^. EDAM is a comprehensive ontology of well-established, familiar concepts that are prevalent within bioinformatics and computational biology. The biotoolsSchema is a formalized schema (XSD) that also includes EDAM ontology and is used by the ELIXIR Tools & Data Services Registry bio.tools^43^. Bioschemas is a community project built on top of schema.org, aiming to improve interoperability in Life Sciences so resources can better communicate and work together by using a common markup on their websites. Bioschemas has overlap with CodeMeta. These schemas and ontologies do not have formal file formats for inclusion in research software and are rather suited for repositories and registries hosting software metadata.

The suggestions above are tailored toward machine-friendly metadata files. Software metadata in a human-friendly format, i.e. documentation, is also required to comply with the FAIR4RS principles. Suggestions for such documentation are provided in 9 of the reviewed resources with one study fully dedicated to best practices for documenting scientific software^30^. Including a README file (called README, README.txt, or README.md) is the method suggested by all of the resources for maintaining documentation^14, 19, 21, 24, 28, 30, 33, 44^. For more visibility, one of the reviewed resources^24^ suggests maintaining a website using GitHub pages (https://pages.github.com) or Read the Docs (https://readthedocs.org). Several resources are suggested in the reviewed literature to help with preparing documentation for research software: Write the Docs, Doxygen, Sphinx3, Javadoc (Java code), Roxygen (R code). Documenting changes between versions of software in a CHANGELOG file is also suggested^19^.

There is clearly a lack of community agreement on metadata files and ontologies to use for documenting research software. Therefore, we used findings from this review, combined with our own assessment and external suggestions we received when consensus was lacking, to establish our recommendations for fulfilling the instructions from Category 2. We recommend including both a codemeta.json and CITATION.cff metadata files in the root directory of the software code. While these two metadata files include some overlapping fields, both are recommended since the codemeta.json file provides a machine-oriented format while the CITATION.cff provides a more human-readable format. Moreover, they both fulfill the requirement of using controlled vocabulary and include typically suggested metadata fields in the FAIR4RS principles. The CodeMeta generator and CFF file initializer are available to help prepare both. We suggest providing all available fields in both files. To align with the FAIR4RS principles, we recommend providing at least the following fields in the codemeta.json file:

- Software name (“name”)
- Software description/abstract (“description”)
- Unique identifier (“identifier”)
- Authors (“givenName”, “familyName”) with their organization name (“affiliation”)
- Keywords (“keywords”)
- Programming Language (“programmingLanguage”)
- First and current release date (“dataPublished” and “dateModified”)
- License used (“license”)

Similarly, we recommend providing at least the following fields in the CITATION.cff file:

- Authors (“given-names”, “family-names”) with their organization name (“affiliation”)
- Software description/abstract (“abstract”)
- Unique identifier (“identifiers”)
- Keywords (“keywords”)
- License (“license”)
- Release date (“date-released”)

In addition, our suggestion is to maintain human-friendly documentation at the very least in a README.md or README.txt file located in the root directory of the software. Mature/complex software may require additional, more sophisticated documentation that can be developed e.g. using tools such as GitHub pages or Read the Docs. To comply with the FAIR4RS principles, the following aspects must be documented: overall description of the software (e.g., in an “About” section), high-level dependencies of the software (e.g., Node or Python version), inputs and outputs of the software, parameters and data required to run the software, the standards followed, how to contribute to the software, and how to cite the software. We also recommend following any community-agreed standard documentation approach when available (e.g., the Common Workflow Language (CWL)^45^ for describing command line tools). Finally, we recommend documenting changes between different versions of the software in a file called “CHANGELOG” using plain text or markdown syntax. Locate it in the root directory of the software. We suggest following the “Keep a changelog” (https://keepachangelog.com) conventions for the content of the CHANGELOG file and using the Semantic Versioning v2.0.0 (https://semver.org) for software version numbers.

#### Category 3: Provide a license

From the reviewed resources, 18 have made suggestions about a suitable license for research software^6, 10, 12–15, 18, 21–24, 26, 31, 33, 36, 46, 47^. All agree that it is preferable to use an open-source license to make the software as reusable as possible. Since there are a large variety of open-source licenses available, it should be possible to find one that fits everyone’s needs^13^. Therefore, if an open-source license is not used, that decision is expected to be properly motivated^31^. Most of the reviewed resources don’t explicitly suggest a specific license, but typically encourage choosing a license approved by the Open Source Initiative (OSI) (https://opensource.org/licenses) since they are well known and understood thus making reuse easier^18, 24^. Some of the reviewed resources do recommend using permissive licenses since they have very few restrictions making them optimal for reuse^6, 21^. Some explicitly encourage the use of the permissive MIT and Apache 2.0 licenses^12, 14, 18, 21, 26, 47^. The Software Package Data Exchange (SPDX) (https://spdx.dev), Choose a License (https://choosealicense.com), and the lesson on license from the 4 Simple Recommendations for Open Source Software are some suggested resources for getting help with selecting a suitable license. It is typically suggested to include the license terms in a LICENSE.txt or LICENSE.md file stored in the root directory of the software^21, 22, 24, 28^.

Based on these findings, combined with our own assessment and external suggestions we received, our recommendation for fulfilling the instructions from Category 3 is to include the license term in a LICENSE.txt or LICENSE.md file located in the root directory of the software. While the FAIR4RS principles do not require research software to be open-source, we highly recommend using a license approved by the Open Source Initiative (OSI). Amongst those licenses, we encourage the use of the permissive MIT or Apache 2.0 licenses. Resources such as Choose a License and/or the SPDX License List can be used to help with selecting a license and including standard terms in the LICENSE file. It is suitable to select the license at the beginning of the development of the software so that it is easier to ensure that the software dependencies are compatible with the license.

#### Category 4: Share software in a repository

From the reviewed resources, 9 have made a recommendation on the files to share for software. Sharing the source code is recommended by most of them to optimize reusability^8, 9, 11, 23, 29, 48^. Providing an executable^9^ and some input/result data^19, 23, 48^ when available is also recommended.

From the reviewed resources, 27 made a suggestion about repositories to use for sharing research software. Three types of repositories are typically mentioned: archival, deployment, and domain-specific. The Registry of Research Data Repositories (re3data.org) is often mentioned as a tool for finding a suitable repository.

Zenodo (https://zenodo.org) is the most commonly suggested archival repository across all reviewed studies^6, 8, 12–15, 17–19, 21, 25–28, 31, 32, 34, 36, 47–50^. Funded by the European Commission and developed and hosted by the Centre Européen pour la Recherche Nucléaire (CERN), Zenodo allows any digital resources, including research software, to be shared and preserved in line with the FAIR principles (https://about.zenodo.org/principles). Zenodo assigns a separate Digital Object Identifier (DOI) for each released version of a software and also creates a high-level DOI that refers to all versions^51^. Of the 50,000 plus DOIs registered for software, more than 80% were registered via Zenodo showing its popularity amongst the research software community^32, 52^. Archiving from GitHub is made easy since it is possible to integrate GitHub and Zenodo such that each GitHub release of a software is automatically archived on Zenodo. Figshare (https://figshare.com) is also a highly recommended archival repository^6, 14, 15, 18, 21, 23, 27, 48–50^. Figshare is a repository supported by Digital Science where researchers can preserve and share any research outputs, including research software, in line with the FAIR principles^53^. Archiving GitHub repositories on Figshare is also made easy through a tool developed in collaboration with the Mozilla Science Lab (https://mozillascience.github.io/). Software Heritage (https://www.softwareheritage.org) is a commonly suggested archival repository as well^6, 8, 12–14, 17, 19, 25, 31, 32, 50, 54^. Software Heritage is an international initiative to provide a universal archive and reference system for all software. It automatically and regularly harvests source code from version control system platforms such as GitHub and anyone can also initiate the submission of their source code for archival. Software Heritage assigns intrinsic identifiers that follow a standardized format called SoftWare Heritage persistent IDentifiers (SWHIDs). Contrary to Zenodo and Figshare, unique identifiers are assigned to all artifacts over all levels of granularity such as project status, project release, state of source code, and code fragment, which can be useful to unambiguously refer to any component of a software.

Some of the reviewed resources^6, 17, 18, 25, 31, 50^ mention using deployment repositories which are typically language-specific such as PyPI (https://pypi.org) and Conda (https://conda.io) for Python packages, CRAN (https://cran.r-project.org) for R packages^55^, and Dockstore (https://dockstore.org) for Docker-based tools. Only two biomedical-specific repositories were mentioned in the reviewed studies^6, 48^. ModelDB for computational neuroscience models^56^ and Bioconductor for R-packages aimed at the analysis of genomics data^57, 58^.

Based on these findings, combined with our own assessment and external suggestions we received, our recommendations for fulfilling the instructions from Category 4 are threefold. If applicable, we suggest sharing each version of a software on a deployment repository (e.g., PyPI or Conda for Python packages, CRAN for R package, Dockstore for Docker-based tools). While such repositories do not provide unique and persistent identifiers as required in the FAIR4RS, this is still useful for increasing the findability and reusability of the software. In addition, we suggest always archiving each version of a software on Zenodo or Figshare as they both allow to archive software in line with the FAIR4RS principles. Zenodo is preferable due to its popularity for archiving research software. The source code of the software with all the above-mentioned metadata files must be archived. Executables and sample input and out data, if available, must be archived as well. Finally, we suggest archiving the software on Software Heritage directly from your version control system platform as it will automatically assign a unique identifier for all levels of granularity. Note that there is no community agreement on the identifier (e.g., DOI or SWHID) to use for software, which is a gap that will need to be addressed by the community.

#### Category 5: Register on a registry

From the reviewed resources, 8 made a suggestion about registries to use for research software^6, 8, 12, 17, 24, 25, 34, 59^. The repositories Zenodo, Figshare, Software Heritage, CRAN, PyPI, and Conda are also mentioned as registries in the reviewed literature since they require specific metadata that is used for indexing. bio.tools (https://bio.tools) is the most commonly suggested registry. Supported by ELIXIR, the European Infrastructure for Biological Information, bio.tools provides a registry of biomedical tools^43^ and contains more than 28,000 registered tools (as of June 2023). As explained in the metadata files section above, bio.tools uses the biotoolsSchema, including EDAM ontology, to describe registered bioinformatics software. A unique ID called biotoolsID is assigned to each software. Another possibility, not found in the reviewed resources but known to the authors, is to register the software on the Research Resource Identifiers (RRIDs) portal (https://www.rrids.org) and obtain an RRID. RRIDs are unique numbers assigned to help researchers cite key resources, including software projects, in the biomedical literature to improve the transparency of research methods^60^. Note that an RRID identifies a software as a whole, but not its different versions. Moreover, there exist a collaboration between bio.tools and the RRID portal to share software metadata and cross-link entries.

Based on these findings, combined with our own assessment, and external suggestions we received, our recommendation for fulfilling instructions from Category 5 is to register the software on bio.tools. Even if Zenodo and Figshare already act as registries, it is still suggested to register on bio.tools to increase findability and create additional rich metadata about the software following biotoolsSchema. The software can optionally be registered on the RRIDs portal as well. Registering on bio.tools and the RRID portal is only required once but the registry-specific metadata must be updated as needed for each new software version.

### FAIR Biomedical Research Software guidelines

To make the above-mentioned recommendations easy to implement, we organized them into step-by-step guidelines that align with the typical software development process. This led to the FAIR Biomedical Research Software (FAIR-BioRS) guidelines. The first versions of the guidelines (v1.0.0 and v1.0.1) were based on the FAIR4RS principles v0.3. They then evolved as we aligned our work with the FAIR4RS principles v1.0 as presented in the manuscript to the FAIR-BioRS guidelines v2.0.0 presented in **Table 2**. Implementing these guidelines will ensure compliance with the FAIR4RS principles as shown in the crosswalk presented in **Table 3**. We have established a “FAIR-BioRS” organization on GitHub to maintain all the related resources (http://github.com/FAIR-BioRS). These guidelines and the crosswalk are maintained in a GitHub repository called “Guidelines” within that organization. It documents all the versions and is also archived on Zenodo every time a new version is released^61^. This repository will serve as a tool to collect community feedback on the FAIR-BioRS guidelines. New versions of the guidelines, that will be established based on community feedback or evolution of current standards, will be maintained there. We recommend readers to consult it to access the latest versions of the guidelines.

**Table 2.**
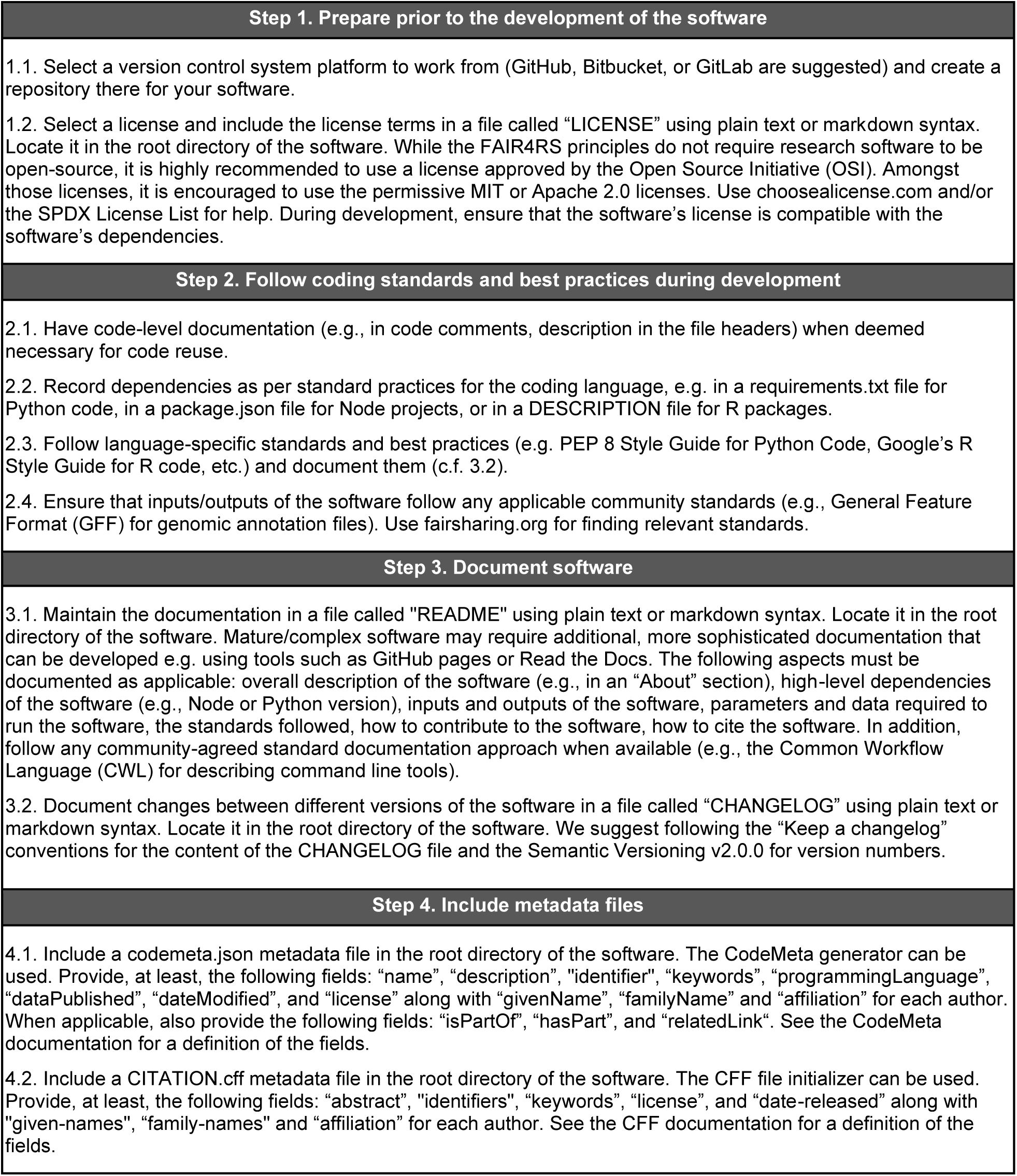
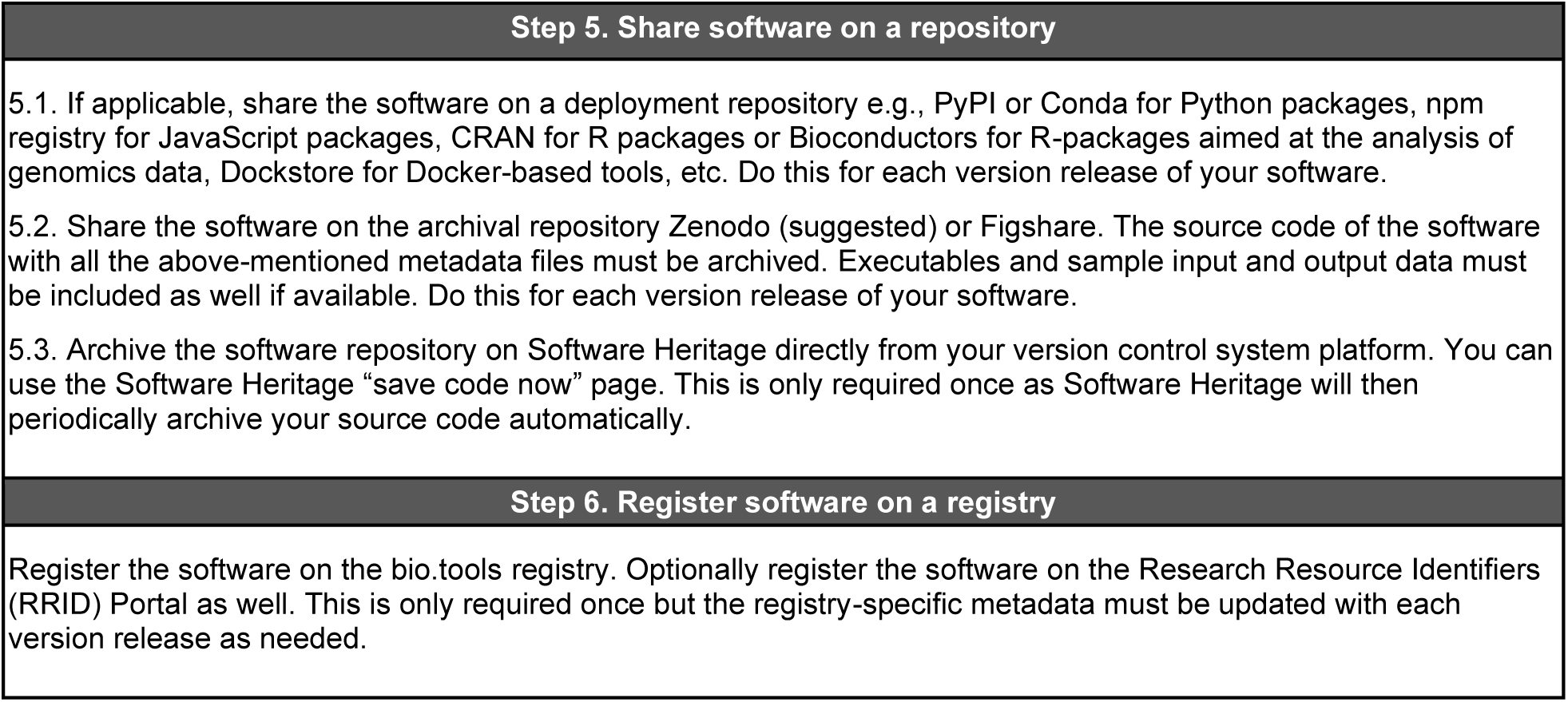
The FAIR-BioRS guidelines version 2.0.0. These guidelines provide a step-by-step process along with reference to relevant resources and tools for making biomedical research software FAIR according to the FAIR4RS principles v1.0. We refer readers to the GitHub repository associated with these guidelines (c.f. Results section) for a markdown version with hyperlinks to relevant resources. New versions of the guidelines, if any, will also be maintained there.

**Table 3.**
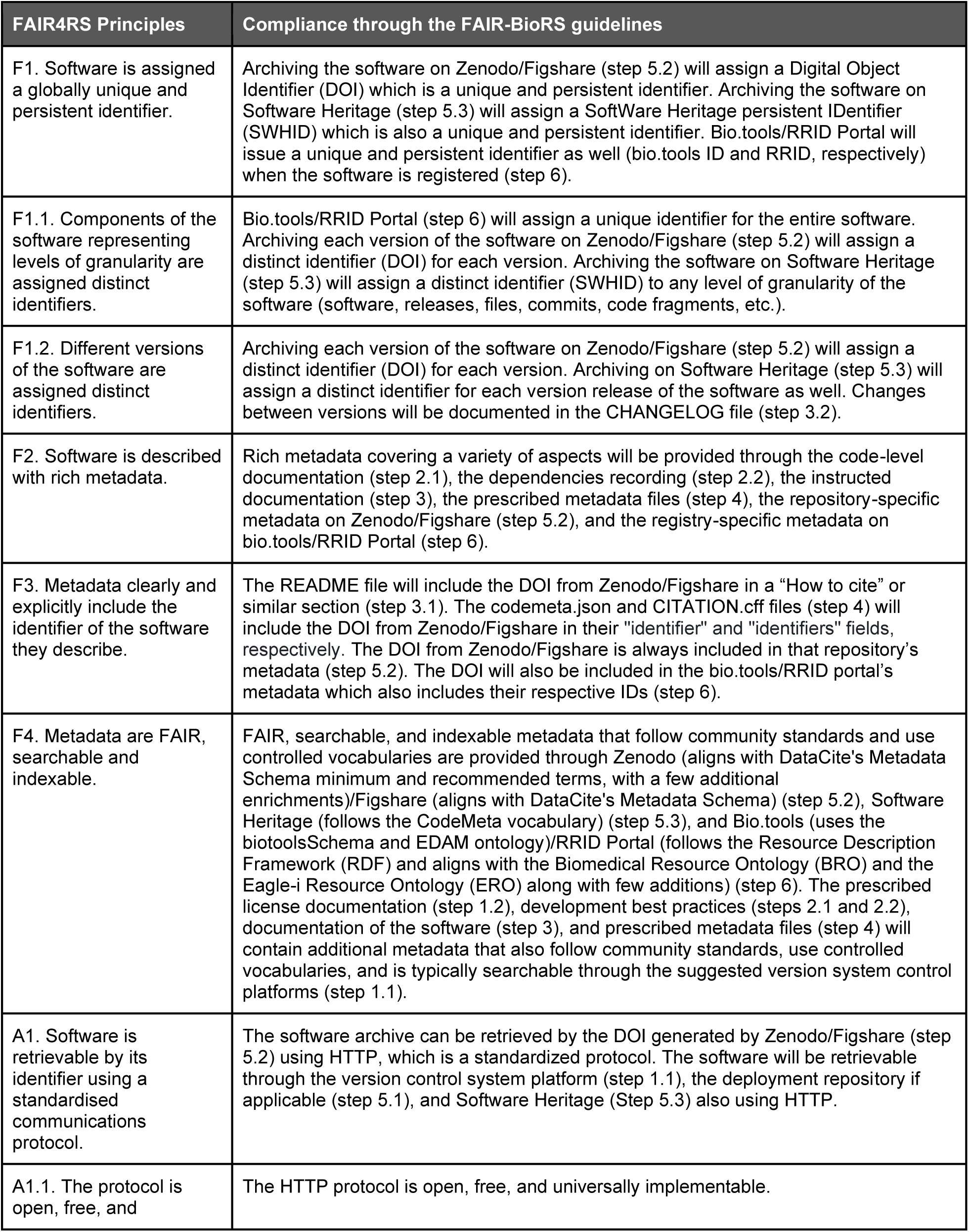
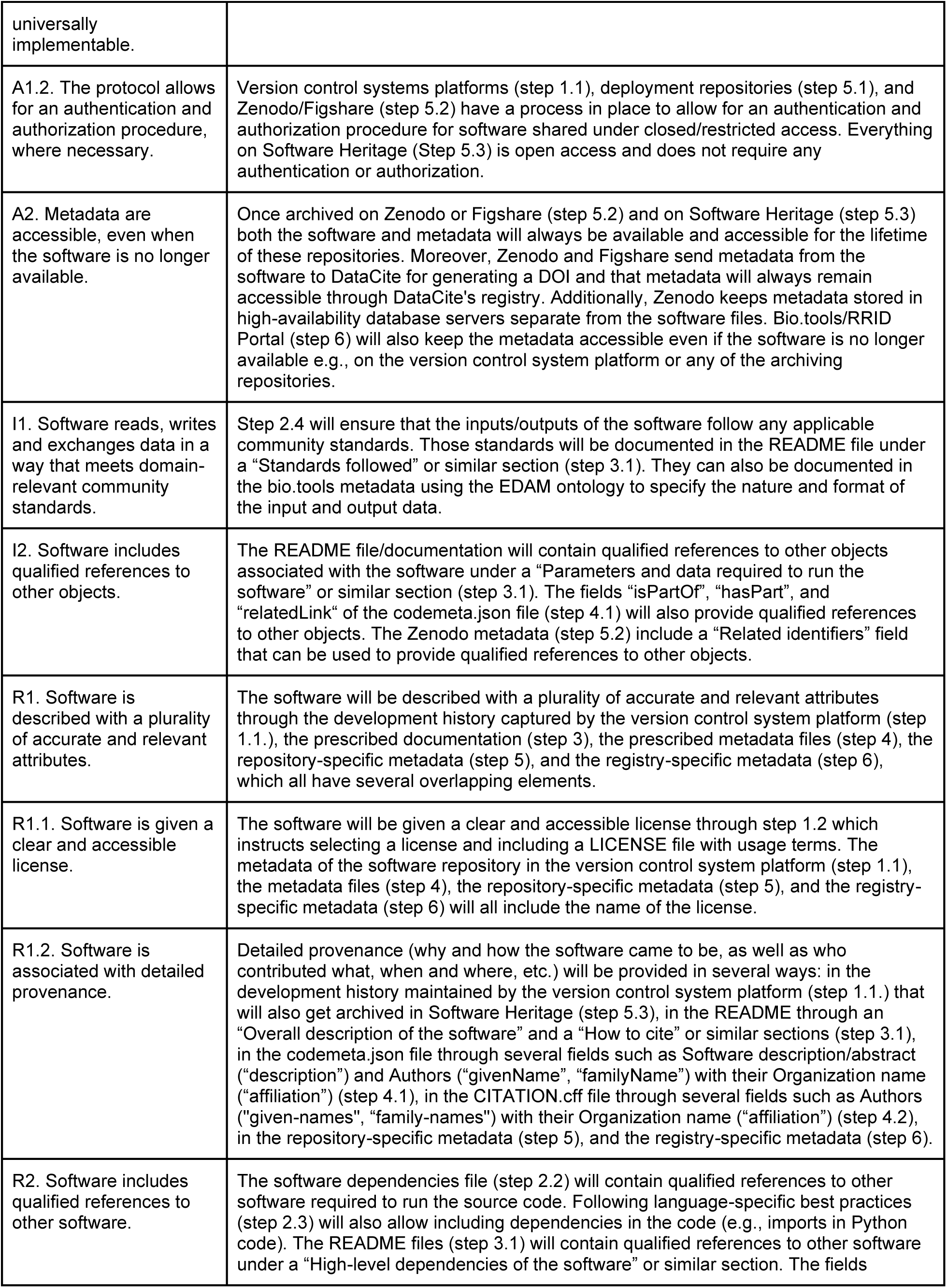
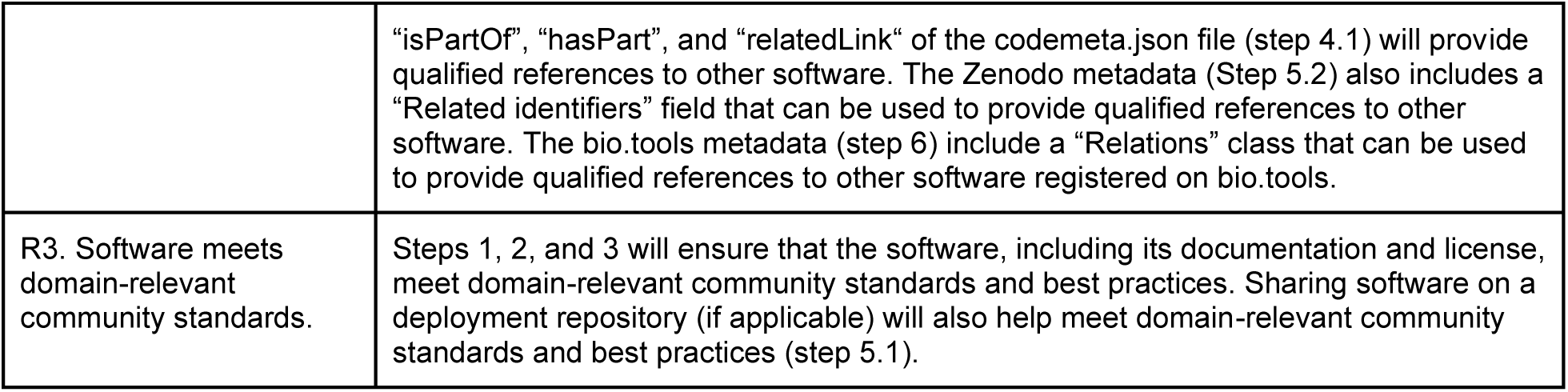
Crosswalk table that explains how the FAIR-BioRS guidelines version 2.0.0 allow complying with the FAIR4RS principles v1.0.

### FAIRshare

Guidelines typically have little effect and are very unlikely to be adopted without proper tools to support them. Tools and resources are available to implement some steps of the FAIR-BioRS guidelines as mentioned in our recommendations above, but they are dispersed and not logically connected to streamline the process. Therefore, we developed and integrated a workflow to implement the FAIR-BioRS guidelines in our software application called FAIRshare. FAIRshare is aimed at streamlining the processes for making biomedical research data and software FAIR. Specifically, FAIRshare combines intuitive user interfaces, automation, and user support resources into a single location to guide and assist the researchers through a suitable workflow for making their research data and software FAIR and sharing it on an adequate repository. FAIRshare is free and developed under the permissible open-source MIT license to encourage community contributions for updating existing workflows and including new ones. Details about the development of FAIRshare are provided in the Methods section. The current version of FAIRshare (v2.1.0) and its documentation (v5.0.0) can be accessed through their respective GitHub repositories and their Zenodo archive (c.f. Code Availability section).

Typically, a user will use FAIRshare to comply with the FAIR-BioRS guidelines when they are ready to publish a new version of their software. The workflow to make research software FAIR with FAIRshare guides users through the steps of the FAIR-BioRS guidelines in a slightly different order to optimize the workflow, as illustrated in **Figure 4**. In step 1, FAIRshare asks the user to select the location of their software-related files. FAIRshare provides users with the option to select files from their computer or select one of their GitHub repositories. If the user selects GitHub, FAIRshare provides an interface to connect FAIRshare to their GitHub account. In step 2, FAIRshare asks a series of “Yes/No” questions to verify that the user has followed applicable standards and best practices during the development and documentation of the software as dictated by Steps 2 and 3 of the FAIR-BioRS guidelines. Links to the relevant resources on standards and best practices suggested by the FAIR-BioRS guidelines are provided in the interface. In step 3, FAIRshare provides a convenient form-type interface so the user can provide information about their software. The fields of the form correspond to fields from the codemeta.json and CITATION.cff metadata files and the user’s entries are used to automatically generate and include these metadata files in the user’s high-level dataset directory. If a codemeta.json file is found at the location selected by the user in step 1, information from that file is automatically pulled by FAIRshare to pre-populate the form. If no codemeda.json file is found and the user files are located in a GitHub repository, FAIRshare will automatically pull available information from the repository metadata (authors, keywords, etc.) and pre-populate related fields of the form. Throughout the forms, tooltips have been included to help users enter the required information. Suggestions are also made for some of the fields to help the user provide inputs rapidly (e.g., “National Institutes of Health” is suggested for the “Funder” field, “Scientific” is suggested for the “Software type” field, etc.). If any of the fields require an entry following a controlled vocabulary for the codemeta.json or CITATION.cff metadata files, FAIRshare typically provides a dropdown list of standardized options to select from to ensure metadata files are generated without error. FAIRshare also makes it mandatory to provide inputs for the metadata fields that were deemed essential by the FAIR-BioRS guidelines. In step 4, the user is prompted to select a license for their software if a LICENSE file is not found in their software file. A dropdown list is provided to select one of the licenses recommended by the OSI. The “MIT” and “Apache 2.0” licenses are suggested in the user interface since they have been deemed optimal for making research software FAIR. The user can read and edit the terms of a selected license directly in the app and can request FAIRshare to include a LICENSE file with associated terms in their software. In step 5, the user is prompted to select an archival repository to share their software files. In the current release, FAIRshare supports sharing on both Zenodo and Figshare. A convenient interface is provided to connect FAIRshare to the user’s Zenodo or Figshare account using a token. Upon login, the user is prompted to provide information about the software that is required by the selected repository. FAIRshare provides a form-type interface such that the user can easily input that information. The interface closely mimics the interface from the selected repository platform to maintain familiarity. Since most of the information required by Zenodo and Figshare is similar to that required in the codemeta.json file, FAIRshare pre-populates the form based on user inputs during step 3. Step 6 depends on the location of the user’s software files as specified during step 1:

- If the files are located on the user’s computer, a summary of the user’s dataset files, including files to be generated by FAIRshare, is provided to the user in step 6. Once the user hits the “Start upload” button, FAIRshare creates a draft deposition on the selected archival repository to reserve a DOI, generates the codemeta.json and CITATION metadata files (with the reserved DOI included) and LICENSE file in the dataset folder, zips the dataset folder, and uploads it in the draft deposition.
- If the files are located on a GitHub repository, FAIRshare provides an interface to include any additional files in their archive if desired besides those already on their GitHub repository (e.g., executables). Then, a summary of the user’s dataset files, including files to be generated by FAIRshare, is provided to the user in step 6. Once the user hits the “Start upload” button, FAIRshare creates a draft deposition on the selected archival repository to reserve a DOI, generate and push the codemeta.json and CITATION metadata files (with the reserved DOI included) and LICENSE file on their GitHub repository. The GitHub repository content is then downloaded, and any additional files specified by the user are included in the downloaded folder. FAIRshare then zips that dataset folder and uploads it into the draft deposition.

**Figure 3.**
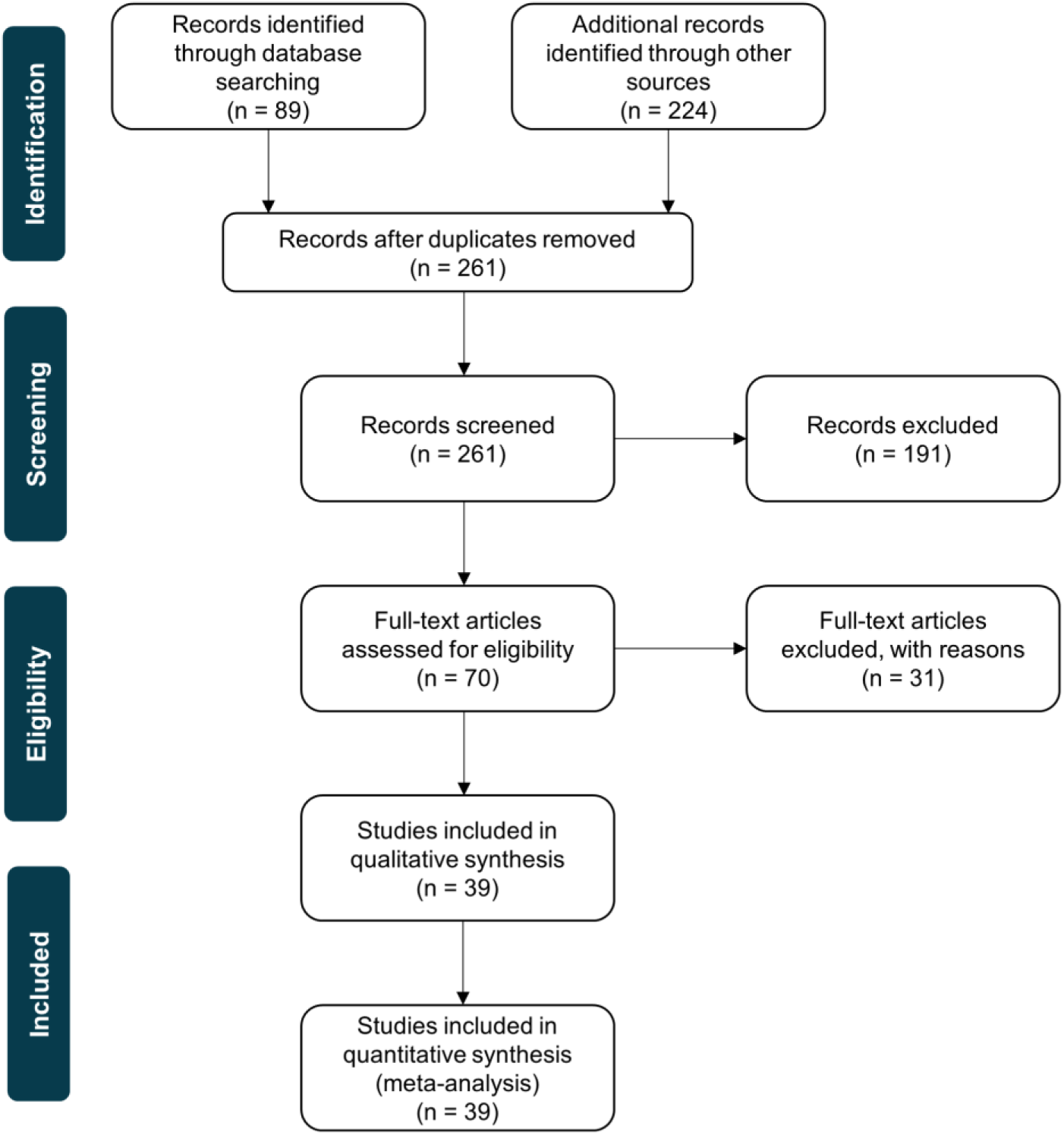
PRISMA (Preferred Reporting Items for Systematic Reviews and Meta-Analyses) diagram providing an overview of our literature review process.

**Figure 4.**
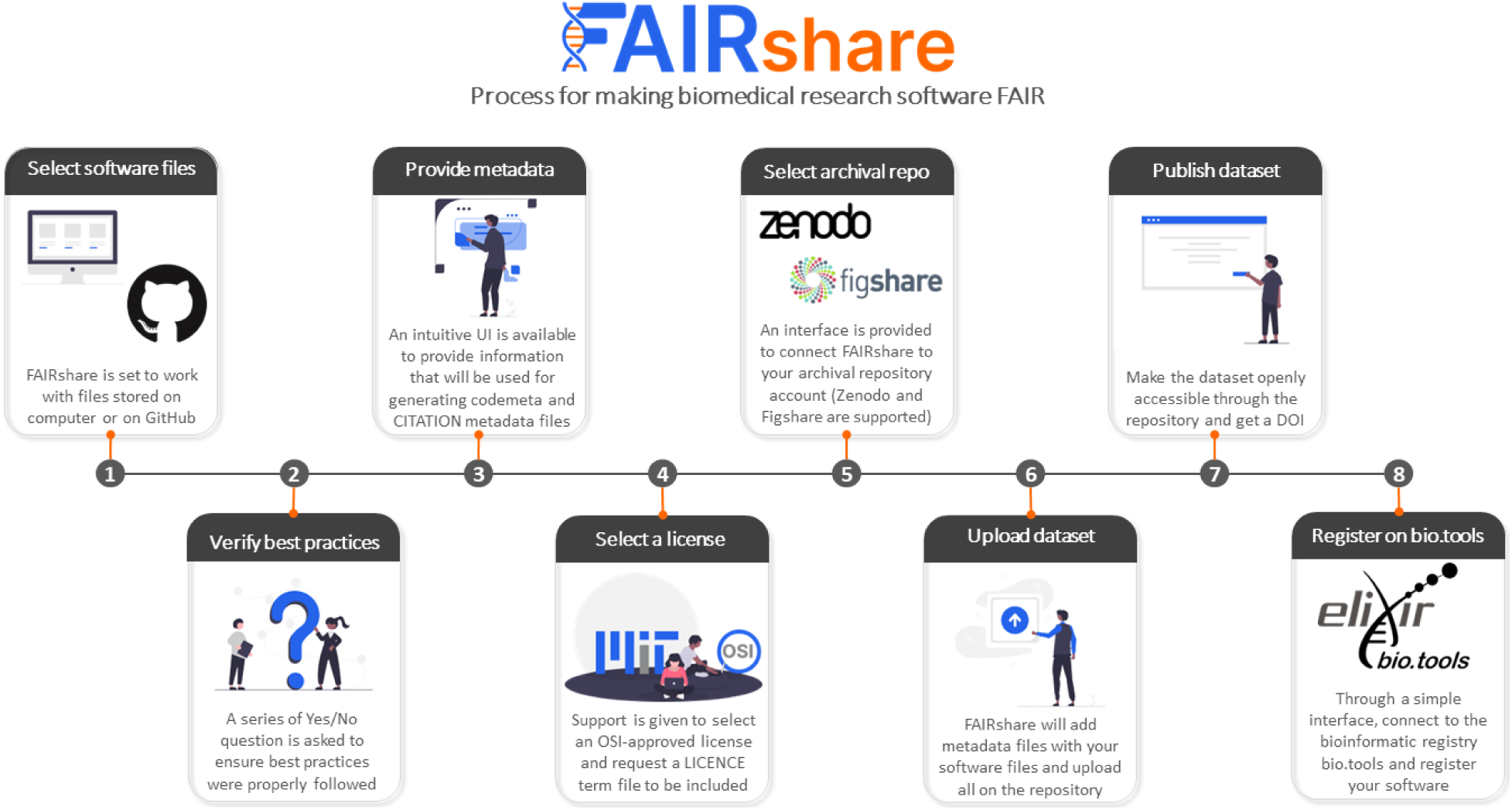
Illustration of the step-by-step guided workflow in FAIRshare for implementing the FAIR-BioRS guidelines and making biomedical research software FAIR. Users of FAIRshare can follow this workflow when they are ready to publish a new version of their biomedical research software.

Upon completion, the user is prompted in step 7 to publish their dataset on the archival repository, such that it becomes publicly accessible and is also given the option to create a GitHub release of the same. A message is shown to the user after completing step 7 to encourage them to archive their software on Software Heritage and register it on bio.tools. If the user selects to register on their software on bio.tools, an interface is provided in step 8 where the user can connect FAIRshare to their bio.tools account, enter basic metadata required by bio.tools (which is prepopulated based on information already available from the codemeta.json file), and register their software.

Besides combining all required resources for making software FAIR into a single interface, FAIRshare also provides some advantages over existing tools. For instance, the Zenodo/GitHub integration allows users to automatically create a Zenodo deposit every time a GitHub release is created. However, it does not allow to get the DOI of the deposit before it is published and therefore it cannot be included in the codemeta.json and CITATION.cff metadata files, which violates principles F3 that prescribes for metadata to “include the identifier of the software they describe”. This is addressed in FAIRshare as it creates a draft deposit of the software first on Zenodo and Figshare to reserve a DOI and include it in the metadata files before the software is published on either archival repository.

## Discussion

While research software constitutes the backbone of biomedical research, the amount of effort dedicated to ensuring software reusability and long-term sustainability is nowhere near the effort dedicated to data. The Findable, Accessible, Interoperable, and Reusable (FAIR) guiding principles published provide a foundation for managing digital research objects, including software, such that their reusability by humans and machines is optimized. However, they fail to capture the specific traits of software such as dependencies and versioning. The Research Data Alliance (RDA) FAIR for Research Software (FAIR4RS) Working Group derived much-needed reformulated FAIR principles to address this shortcoming. Just like the original FAIR guiding principles, the FAIR4RS guiding principles are still aspirational and do not provide practical instructions and actionable items to the researchers since they may be domain-specific while the principles are meant to cover all research software. To fill this gap, we proposed in this work the first minimal and actionable step-by-step guidelines for biomedical researchers to make their research software compliant with the FAIR4RS principles. Our process for deriving these guidelines started with establishing high-level instructions to fulfill each of the FAIR4RS principles based on FAIR4RS v1.0 publications. Noticing common themes across the instructions, we organized them into five high-level categories. For each category, we identified outstanding questions we needed to answer for fulfilling the associated instructions. Subsequently, we conducted a thorough literature review to find out available practices for answering these questions. Through analysis of our findings and our own assessment, we ultimately established the FAIR-BioRS guidelines, which are the first minimal and actional guidelines for making research software FAIR by fulfilling all the requirements of the FAIR4RS principles.

The reviewed resources included the best practices suggested by the NIH for sharing research software (https://datascience.nih.gov/tools-and-analytics/best-practices-for-sharing-research-software-faq). Therefore, the FAIR-BioRS guidelines align with the recommendations of the NIH for sharing research software. Additionally, the current version 2.0.0 of the FAIR-BioRS guidelines incorporate suggestions from various communities. Indeed, after establishing v1.0.1 of the guidelines, we initiated a significant effort to raise awareness about the guidelines and get community feedback. The guidelines were presented^62^ at the Bioinformatics Open Source Conference (BOSC) 2022 organized by the Open Bioinformatics Foundation (OBF). We also introduced the FAIR-BioRS guidelines as a topic of collective work during the CollaborationFest that followed BOSC, which allowed for in-depth discussions with OBF members. The guidelines were also presented to SciCodes, the consortium of scientific software registries and repositories. In addition, the guidelines are being implemented as part of the Software Development Best Practices of the AI-READI project^63^, a large-scale human-data collection and tools development project funded by the NIH Bridge2AI Program. Finally, the guidelines were also presented through various webinars. Overall, we received valuable suggestions on v1.0.1 of the FAIR-BioRS guidelines during those events which helped greatly in shaping v2.0.0 presented in this manuscript. Our community outreach effort is still ongoing. Our abstract on the FAIR-BioRS guidelines v2.0.0 has been selected for presentation at BOSC 2023. We are in discussion with the RDA Software Source Code Interest Group, the maintenance home for the FAIR4RS principles, to present the FAIR-BioRS guidelines. We are similarly planning a presentation for the Elixir Tools team. We have also initiated discussions with the NIH to recommend the FAIR-BioRS guidelines in addition to or in lieu of their current guidelines for sharing research software that does not fully align with the FAIR4RS principles. We are hoping such effort will help with raising awareness about the FAIR-BioRS guidelines and lead to their adoption by various biomedical communities.

To ensure the FAIR-BioRS guidelines are easily accessible and widely embraced by researchers, we have developed a user-support tool to streamline the process of implementing the FAIR-BioRS guidelines. This tool is incorporated in FAIRshare, a software application aimed at simplifying the processes for making biomedical research data and software FAIR. FAIRshare takes the user’s software-related dataset as input when they are ready to publish a new software version and walks them step-by-step into implementing the FAIR-BioRS guidelines through an intuitive graphical user interface and automation. This way, researchers can implement the FAIR-BioRS guidelines even without prior knowledge of them and in fact learn about these guidelines along the way. FAIRshare is free and open-source to encourage community contributions to keep up with the ever-evolving FAIR guidelines.

While establishing the FAIR-BioRS guidelines, we identified several practical gaps and needs for complying with the FAIR4RS principles. We established the best possible guidelines to comply with each of the FAIR4RS principles despite these gaps using our own understanding of research software and external feedback when consensus when there was such a gap. Mainly, there is a lack of community agreement on standards and best practices for developing and documenting software. In addition, there is a lack of community agreement on metadata files and ontologies to use for describing biomedical research software. There is a need to consolidate the various developments (CodeMeta, CFF, Bioschemas, etc.) so developers of biomedical research software can easily navigate through them and ideally use a single metadata file in their software. Moreover, there is a lack of community agreements on standard file formats to use for different biomedical data types, which prevent standardization of the data that software read, write, and exchange. While FAIRsharing is a useful registry, it is still very overwhelming to navigate through the large amount of existing standards, and consolidation is required here as well. There is also a lack of agreement on a suitable identifier to use for a software (DOI, SWHID, bio.tools ID, RRID, etc.). We hope that our work will help bring awareness about these needs to the relevant communities. Our work and choices made in the FAIR-BioRS guidelines may even guide them in their efforts to address them.

We expect the FAIR-BioRS guidelines and related resources to evolve over time based on several factors. Given the gaps mentioned above, we made several decisions for suggesting the different guidelines. These decisions may evolve as we receive more suggestions and feedback from the research software community, for instance, to remove some guidelines as they may be above what is expected by the FAIR4RS principles or to add more to cover missing areas (such as linting guidelines, using codes of conduct, community governance, etc). Moreover, the guidelines can evolve as new community agreements are established to address the above-mentioned gaps. We have thus established a dedicated FAIR-BioRS GitHub organization to maintain all related resources. It includes a repository where new versions of the FAIR-BioRS guidelines will be maintained and published. This repository will also be used to receive community feedback and suggestions through the GitHub issues. We plan next to open ownership of the FAIR-BioRS GitHub organization to other members of the biomedical research software community such that this eventually becomes a community-driven effort that is perpetuated over time.

We expect next to establish more specific guidelines for different types of software (e.g., establish the FAIR-BioRS Python package guidelines or the FAIR-BioRS Jupyter notebook guidelines) such that it becomes even easier for developers of biomedical research software to comply with the FAIR4RS principles. In parallel, we plan to integrate additional resources in FAIRshare such as sharing on Software Heritage, registering on the RRID portal, or working from other version control system platforms than GitHub such as Bitbucket and GitLab. We also plan to include additional automation to further streamline the FAIRshare process for making research software FAIR. These include for instance automatically validating the implementation of language-specific standards or prefilling keywords and descriptions using Natural Language Processing. A major limitation of FAIRshare is that it is intended to help only when a user is ready to publish a new version of their software, when it might be inconvenient for the user to go back and make changes to their source code, e.g. for aligning with language specific standards or adding in code comments. We are therefore planning the development of a tool that integrates directly with version control system platforms and assists users in complying with the FAIR-BioRS guidelines right from the beginning and throughout the development of their software. As the FAIR-BioRS guidelines, FAIRshare, and other resources related to the FAIR-BioRS guidelines are expected to evolve over time, we have established a dedicated GitHub repository called “Hub” in the FAIR-BioRS organization (https://github.com/FAIR-BioRS/Hub). We refer to that repository to access the latest versions of all resources related to the FAIR-BioRS guidelines and track the interaction between their different versions.

We are hoping the biomedical research software community will embrace FAIR practices by making the FAIR-BioRS guidelines an integral part of their software development practices, which will optimize the reusability of software and ultimately increase the pace of discoveries and innovations in the field.

## Methods

### Definition of research software

The FAIR4RS Working Group was constituted of four subgroups, each focusing on different aspects of their effort. Group 3 focused on defining research software through a thorough review of the literature. In the outcome of their effort, they provided the following short and concise definition: “Research Software includes source code files, algorithms, scripts, computational workflows and executables that were created during the research process or for a research purpose. Software components (e.g., operating systems, libraries, dependencies, packages, scripts, etc.) that are used for research but were not created during or with a clear research intent should be considered software in research and not Research Software”^64^. The actionable guidelines developed in this work are for research software based on this definition (although they may be applicable beyond). This includes code, scripts, models, notebooks, libraries, executables, and other forms as long as it fits the previously cited definition. Given that our work was born with the aim of making COVID-19-related data and research software FAIR, our focus was on deriving guidelines for biomedical research software, although some of the findings may be applicable to other fields of research.

### Deriving concise instructions for fulfilling the FAIR4RS principles

The FAIR4RS principles v1.0 contain 17 guiding principles for FAIR research software (vs 15 in the original FAIR principles), including 6 for Findability, 4 for Accessibility, 2 for Interoperability, and 5 for Reusability. We analyzed details about each principle provided in the FAIR4RS v1.0 write up^11^ and its introduction manuscript^12^. Based on these details, we derived concise instructions to be followed for fulfilling each of the principles (provided in the “fair4rsv1.0Instructions” tab of the “data.xlsx” file, c.f. Data Availability section), i.e., instructions that need to be followed for implementing each principle. We identified that these instructions had overlapping themes and were fitting overall into five actionable categories: 1) Develop software following standards and best practices, 2) Include metadata, 3) Provide a license, 4) Share software on a repository, 5) Register on a registry. The action-based instructions were thus regrouped into these categories into one instruction paragraph per category as shown in **Table 1**. Looking at the instructions for each category, we realized that there were several outstanding questions that needed to be answered for deriving actionable guidelines to fulfill the instructions. These questions are provided in **Table 1**.

### Literature review for practical implementation of FAIR4RS principles

We conducted a literature review to help with answering these questions and identifying actionable recommendations for fulfilling the instructions from each category. Since discussion around FAIR for research software is very recent and not widespread in the literature, we reviewed literature not only related to FAIR research software, but also related to making research software reusable without necessarily a reference to the FAIR principles. Our review was also not restricted to biomedical-related articles for the same reason. Our overall review strategy consisted of including resources in English language from the following groups:

- Group 1: We started by reviewing the six published resources on FAIR principles for research software^6, 8–12^.
- Group 2: Subsequently, we reviewed the resources referenced in studies from group 1 that were deemed relevant based on their title and abstract.
- Group 3: We also reviewed resources listed as of February 2023 in the FAIR4Software reading materials ^(^https://www.rd-alliance.org/group/software-source-code-ig/wiki/fair4software-reading-materials), FAIR4RS Subgroup 4 reading list of new research^65^, and the Zenodo community page of the FAIR4RS Working Group (https://zenodo.org/communities/fair4rs) that were deemed relevant based on their title and abstract and were not already encountered during the previous steps.
- Group 4: Given our focus on biomedical research software, we also searched literature related to FAIR for research software on PubMed. The start period was set to January 2015 since the concept of FAIR data was introduced then, and the end period was February 2023. We read resources that were deemed relevant based on their title and abstract and were not already encountered during the previous steps.
- Group 5: Finally, we included additional studies available as of February 2023 from the authors’ knowledge that were not already read in the previous groups.

Details about the review process are provided in the PRISMA diagram in **Figure 3**. More details about the review strategy, including the PubMed search, are included in the “reviewStrategy” sheet of the “data.xlsx” file associated with this manuscript (c.f. Data Availability section). A list of all the resources encountered during our review is provided in the “resourcesList” sheet of the same file. Initially, we copied word-for-word the relevant information as written in the reviewed resources. This is available in the “resourcesReview” sheet of the “data.xlsx” file associated with this dataset (c.f. Data Availability section). Subsequently, we only retained keywords such as the name of the suggested metadata file, repository, etc. to facilitate further analysis. Information from any of the topics that applied to only specific fields of research other than biomedical were left out. This is available in the “resourcesReviewKeywords” sheet of the same file.

We developed a Jupyter notebook-based script (c.f. Code Availability section) that used scientific packages such as pandas^66, 67^, Matplotlib^68, 69^, and seaborn^70^ to analyze our review data. We used findings from the reviewed resources, combined with our own assessment and external suggestions (c.f. Discussion section) when consensus was lacking in literature, to derive recommendations for fulfilling the instructions from each category. Finally, we organized the recommendations into step-by-step guidelines that follow the typical software development process such that they are easier to implement. This led to the first minimal and actionable step-by-step guidelines for making biomedical research software FAIR such that all the requirements of the FAIR4RS principles are met, i.e., the FAIR-BioRS guidelines.

### User support tool: FAIRshare

FAIRshare is inspired by the software called SODA for SPARC, which has been developed by our team to assist researchers funded by the NIH SPARC (Stimulating Peripheral Activity to Relieve Conditions) program in making their data FAIR according to the SPARC data standards^71–73^. Specifically, FAIRshare combines intuitive user interfaces and automation to guide and assist the researchers through the suitable process for making their research data and software FAIR and sharing it on an adequate repository. FAIRshare is developed as a cross-platform desktop software application, such that the data of the researchers remain on their computer or a storage medium of their choice until they are ready to share it on a suitable archival repository. It is built using Electron, GitHub’s framework for building cross-platform desktop applications using web technologies (HTML, CSS, JavaScript, NodeJS). Vue3 is used as the frontend framework to build intuitive, interactive, and responsive user interfaces. The backend is built using Flask, a micro web framework written in Python. The use of Python for the backend of the application was motivated by the popularity of Python in the biomedical research field and the availability of relevant existing packages and APIs for curating research data and software. More details are available in the dedicated GitHub repository for FAIRshare (c.f. Code Availability section).

## Data Availability

The data associated with this manuscript consists of a data.xlsx file that contains details about the data collected and processed during the review of relevant resources. Since no FAIR guidelines were found for structuring review data, we structured the dataset according to the SPARC Data Structure (SDS), which provides a broad data and metadata structure to organize biomedical research data according to the FAIR principles^72^. The SPARC data curation software SODA for SPARC^71–73^ was used to organize the data and prepare the metadata files. The dataset is maintained in a GitHub repository called “Data” in the FAIR-BioRS GitHub organization and the latest version associated with this manuscript (v3.0.0) is also archived on Zenodo^74^ under the permissible Creative Commons Attribution 4.0 International (CC-BY) license.

## Code Availability

The code associated with this manuscript consists of a “main.ipynb” Jupyter notebook, the source code of FAIRshare, and the source code of the FAIRshare documentation. The “main.ipynb” Jupyter notebook contains the code used to analyze the findings from the review and to conduct other analysis presented in this manuscript (e.g., generate **Figure 1**). This notebook is available in a GitHub repository called “Code” also maintained in the FAIR-BioRS GitHub organization. The dataset associated with this notebook was made FAIR according to the FAIR-BioRS guidelines using FAIRshare v2.1.0^75^, and shared under the permissible MIT license. The latest version associated with this manuscript (v3.0.0) is archived on Zenodo^76^. The source code for FAIRshare is hosted on GitHub (https://github.com/fairdataihub/FAIRshare). The current version of FAIRshare (v2.1.0) discussed in this manuscript was made FAIR using FAIRshare itself, and shared on Zenodo^75^ under the permissible MIT license. The source code for the FAIRshare documentation is maintained on GitHub as well (https://github.com/fairdataihub/FAIRshare-Docs) and the current version (5.0.0) was shared under the permissible MIT license on Zenodo^77^.

## Acknowledgments

This work was supported by grants for the NIH U01AI150741 and NIH SPARC OT2OD030213.

## Author contributions

B. Patel led the conception and design of the FAIR-BioRS guidelines, managed the acquisition/analysis/interpretation of data from the reviewed studies, supervised the development of FAIRshare, and contributed to drafting/revising the manuscript. S. Soundarajan led the development of FAIRshare and contributed to revising the manuscript. Z. Hu and H. Ménager contributed to the conception and design of the FAIR-BioRS guidelines, and revising the manuscript.

## Competing interests

The authors declare no competing interests.

## Notes

https://github.com/FAIR-BioRS

https://doi.org/10.5281/zenodo.8115012

https://doi.org/10.5281/zenodo.8112100

https://doi.org/10.5281/zenodo.8112631

### Competing Interest Statement

The authors have declared no competing interest.

### Summary of Updates

The FAIR-BioRS guidelines were updated to v2.0.0 as per comments from reviewers (this work is under peer-review) and other community feedback we have received along the ways. The manuscript was updated accordingly.

